## Supplementary figures and images for "Systems acclimation to osmotic stress in zygnematophyte cells"

### Extended Data Figure 1

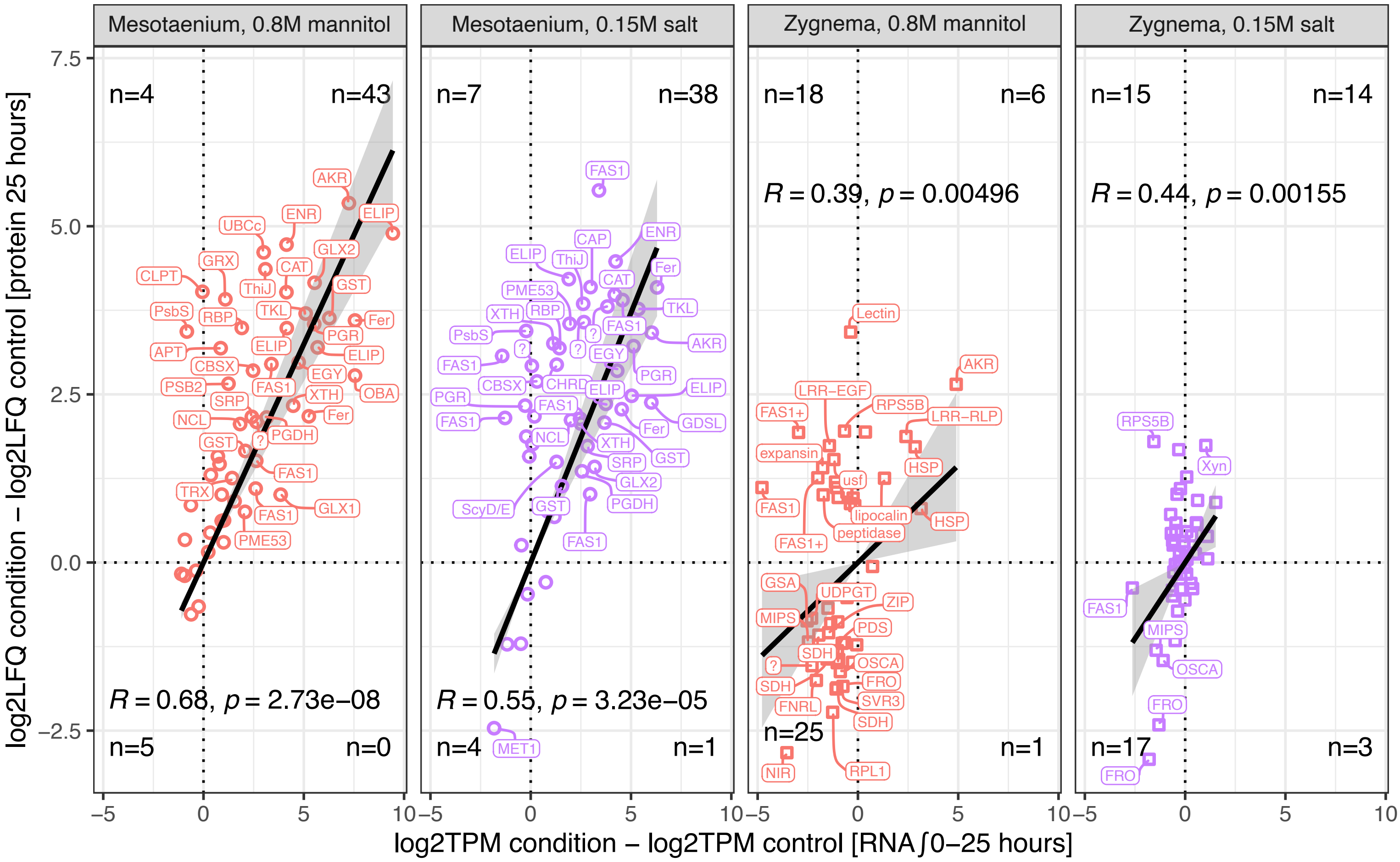
