## Supplementary Figure I for "Systems acclimation to osmotic stress in zygnematophyte cells"

### Supplementary Figure(s) I. new protein groups

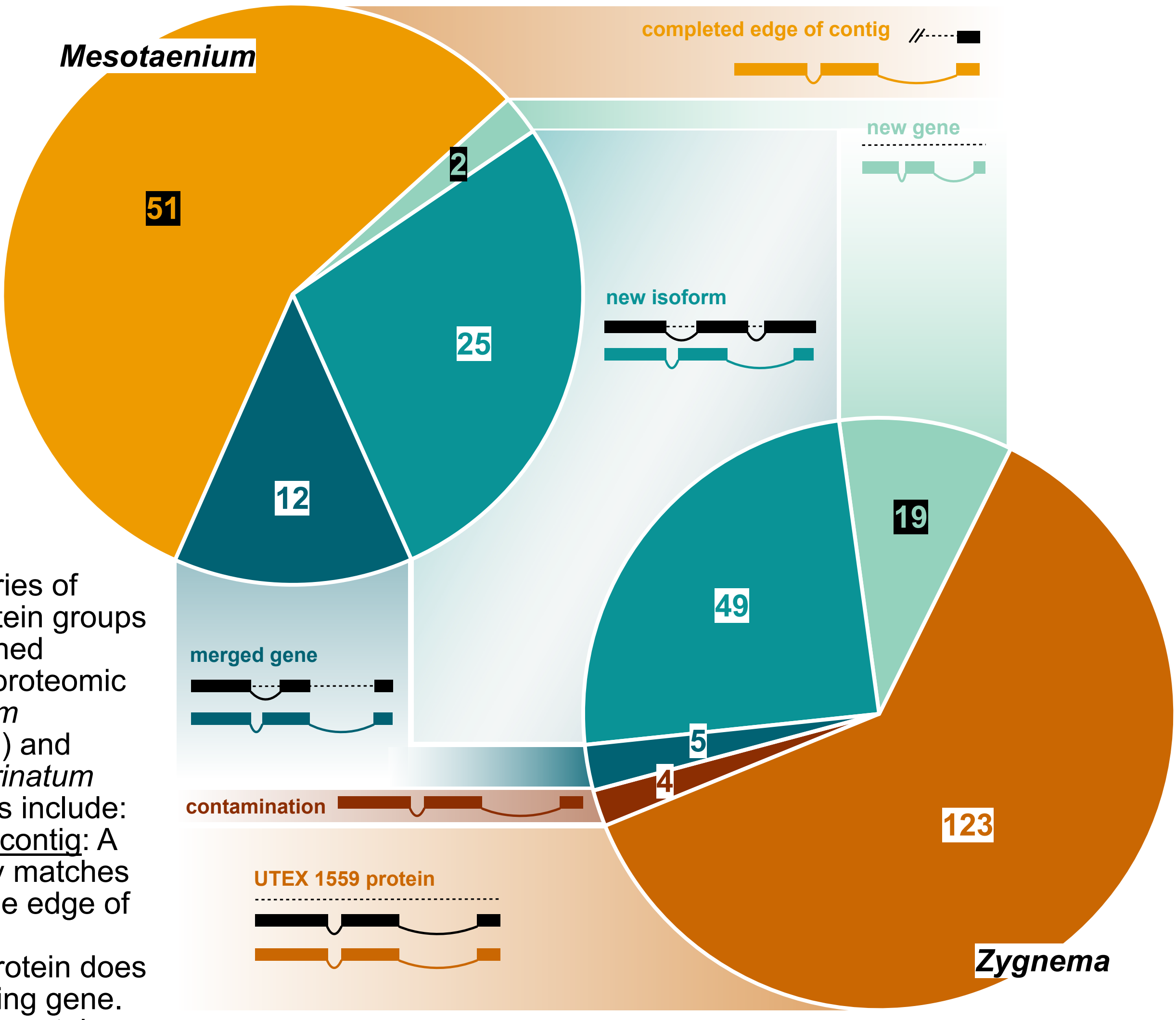


**Figure I.**

**Top Panel:** Categories of newly identified protein groups derived from combined transcriptomic and proteomic data of *Mesotaenium endlicherianum* (Me) and *Zygnema circumcarinatum* (Zci). The categories include:
<u>Completed edge of contig</u>: A new protein partially matches a gene located at the edge of a contig.
<u>New gene</u>: A new protein does not match any existing gene.
<u>New isoform</u>: A new protein group matches an existing gene but represents a novel isoform.
<u>Merged gene</u>: A new protein spans two adjacent genes transcribed in the same direction, effectively merging them.
<u>Contamination</u>: A new protein originates from external contamination, not the studied algae.
<u>UTEX 1559 protein</u>: A new protein is not annotated as a gene in strain 698-1b but is annotated in strain UTEX 1559.

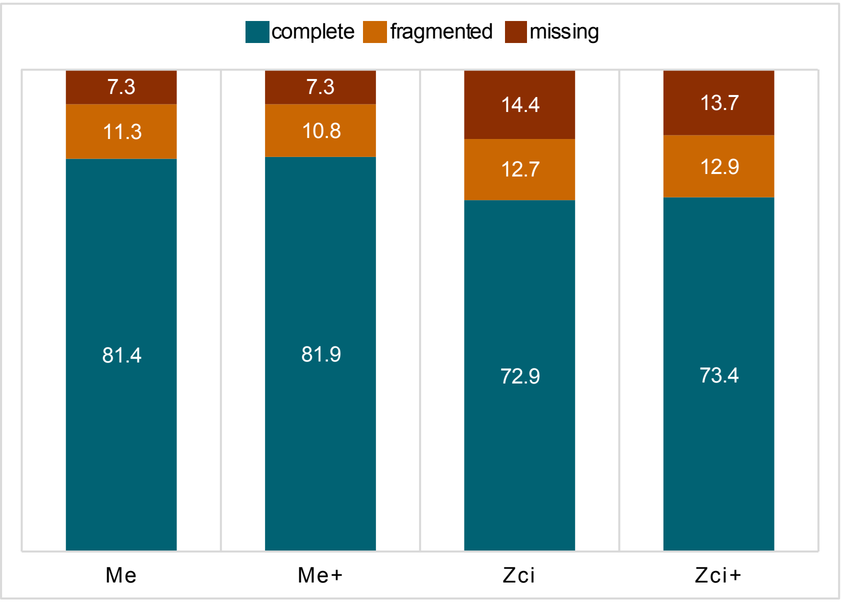


**Bottom Panel:** BUSCO (Benchmarking Universal Single-Copy Orthologs) scores of the proteomes for both algae, shown with (+) and without (-) the newly identified protein groups.
