## Supplementary Figure II for "Systems acclimation to osmotic stress in zygnematophyte cells"

### Supplementary Figure(s) II. Protein-RNA correlations

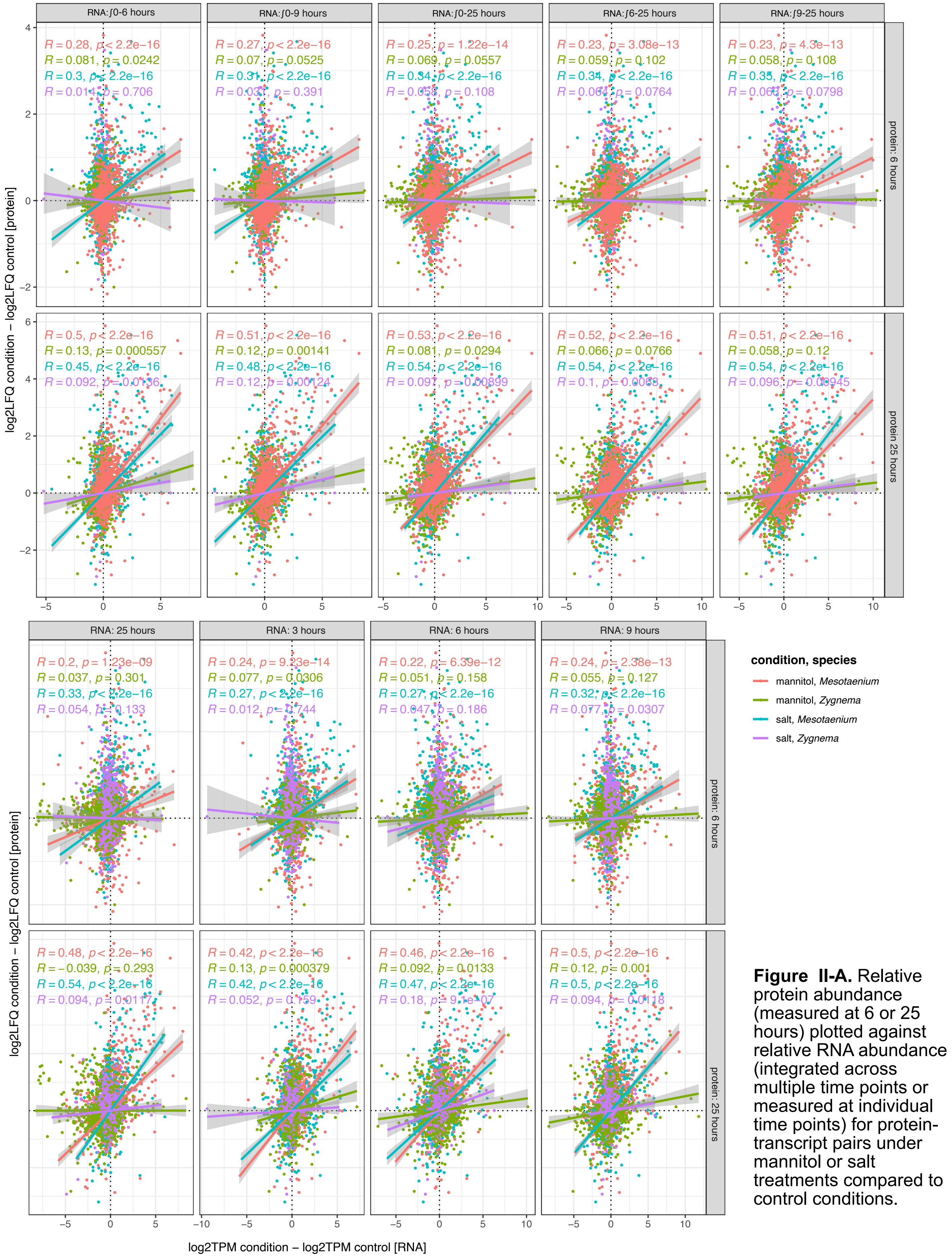


**Figure II-A.** Relative protein abundance (measured at 6 or 25 hours) plotted against relative RNA abundance (integrated across multiple time points or measured at individual time points) for protein-transcript pairs under mannitol or salt treatments compared to control conditions.

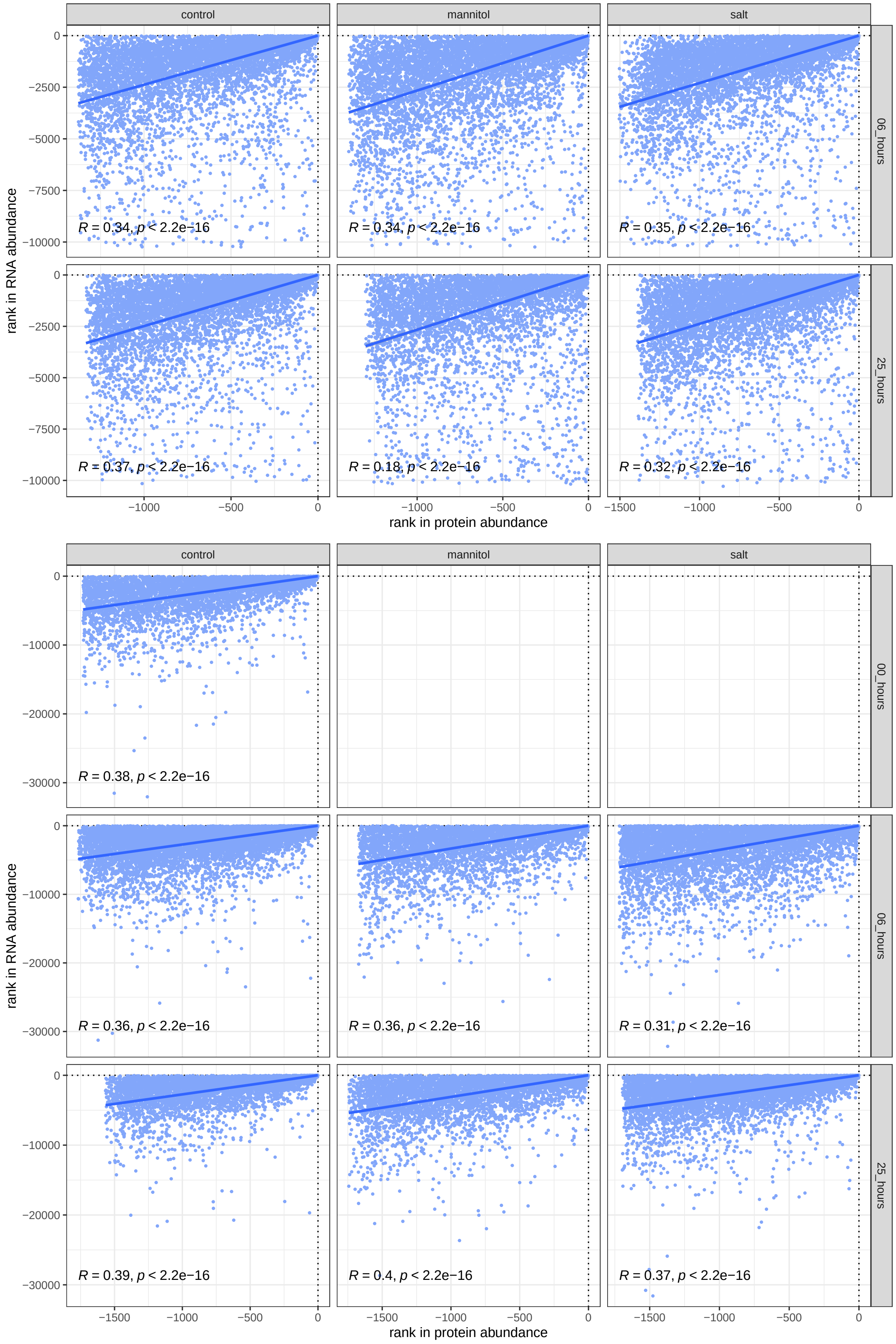


**Figure II-B.** Absolute protein abundance in *Zygnema* (top graph) and *Mesotaenium* (bottom graph) plotted against absolute RNA levels.

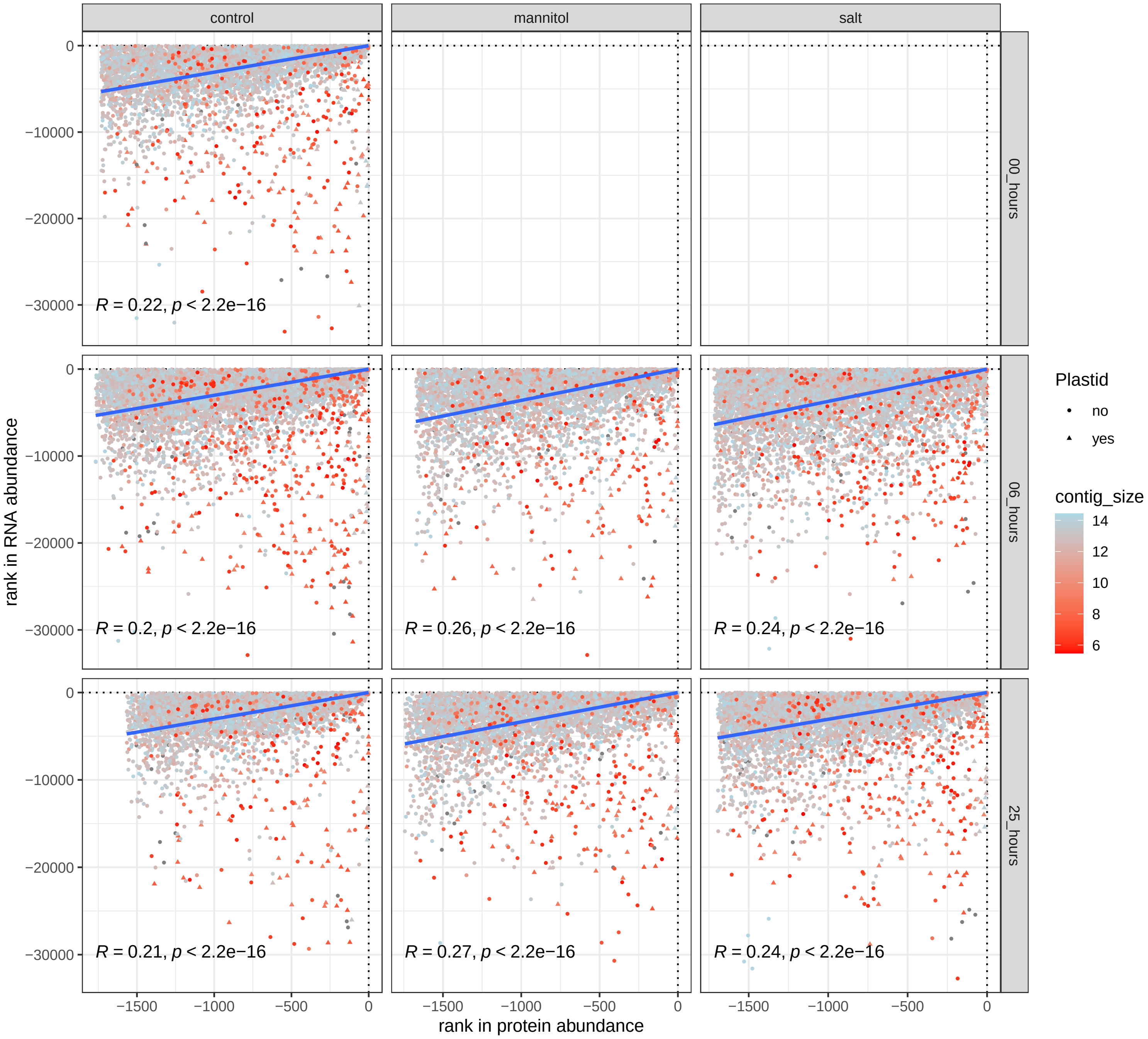


**Figure II-C.** Absolute protein abundance in Mesotaenium (bottom graph) plotted against absolute RNA levels, with data points color-coded based on the size of the contig containing the corresponding protein-coding gene.
