## Supplementary Figure III for "Systems acclimation to osmotic stress in zygnematophyte cells"

### Supplementary Figure(s) III. SWING

**Figure III-A.** Gene regulatory networks generated using Sliding Window Inference for Network Generation (SWING). The large cyan networks display the top 1,750 predicted edges, while the smaller red networks highlight the top 0.1% of predicted edges. In the cyan networks, nodes representing genes with connectivity greater than 12 are labeled with their corresponding transcript.

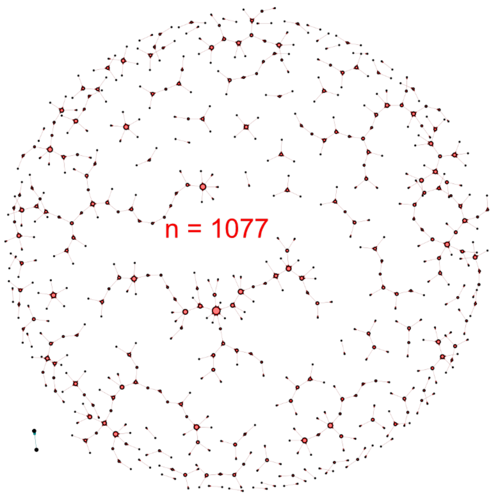


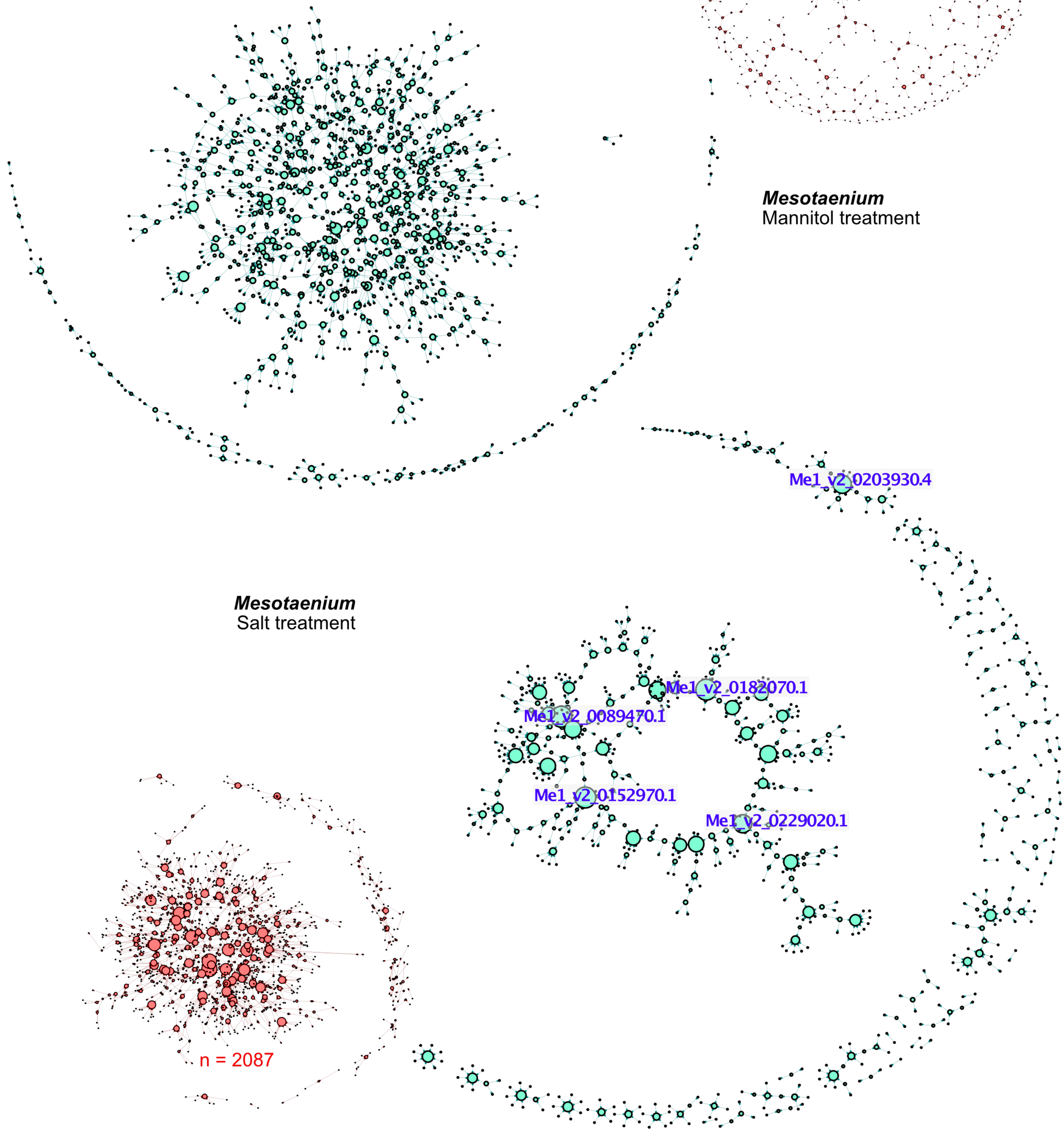

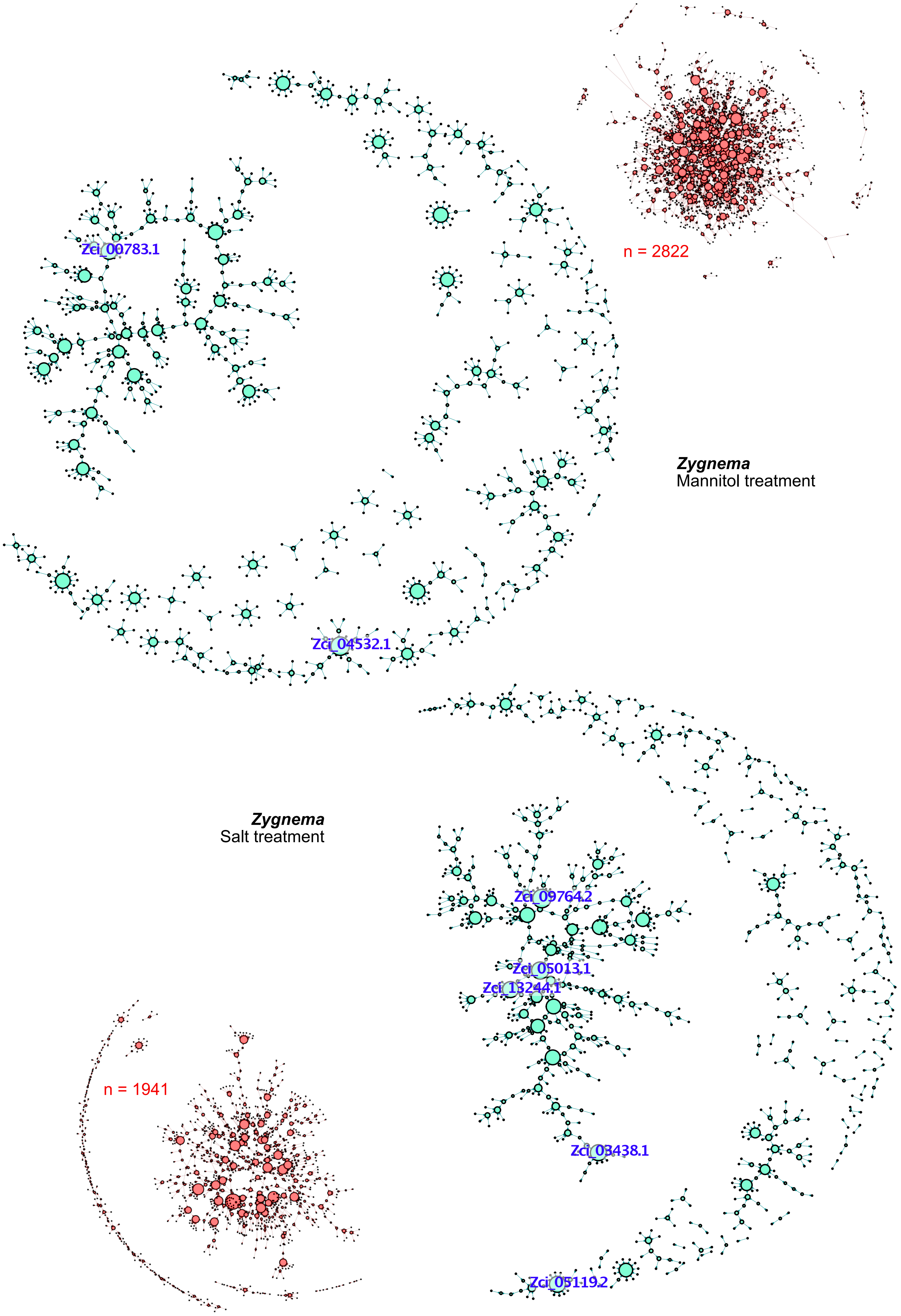

n = 2822
Zci_00783.1
*Zygnema*
Mannitol treatment
Zci_04532.1
*Zygnema*
Salt treatment
Zci_09764.2
Zci_05013.1
Zci_13244.1
n = 1941
Zci_03438.1
Zci_05119.2

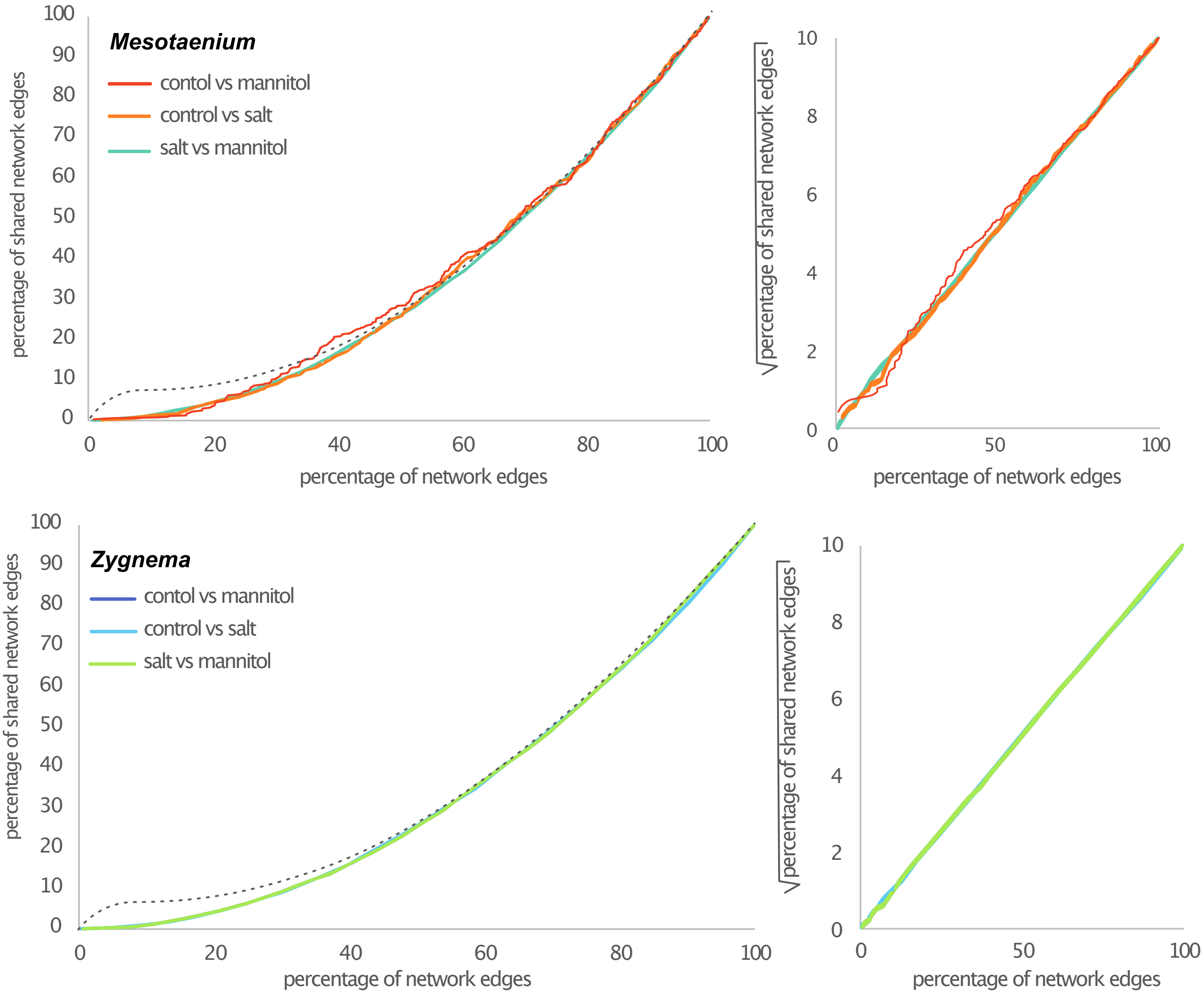


**Figure III-B.** The X-axis represents the percentage of all shared network edges (between two treatments), ranked by score from highest to lowest. The Y-axis shows the percentage of those edges, compared to the total amount of shared edges, that are identical between the two treatments. A steep initial slope in the curve would indicate a higher similarity in the gene regulatory networks between two treatments (as examplified with the dotted line), while the observed trend may suggest uncorrelated gene regulatory networks.
