## Supplementary Figure IV for "Systems acclimation to osmotic stress in zygnematophyte cells"

### Supplementary Figure(s) IV. expression patterns

**Figure IV.** Expression levels (transcripts per million, TPM) for selected genes are presented. Data represent two species: *Zygnema* (Zci) and *Mesotaenium* (Me). Measurements were taken at five time points: 0 hours, 3 hours, 6 hours, 9 hours, and 25 hours.

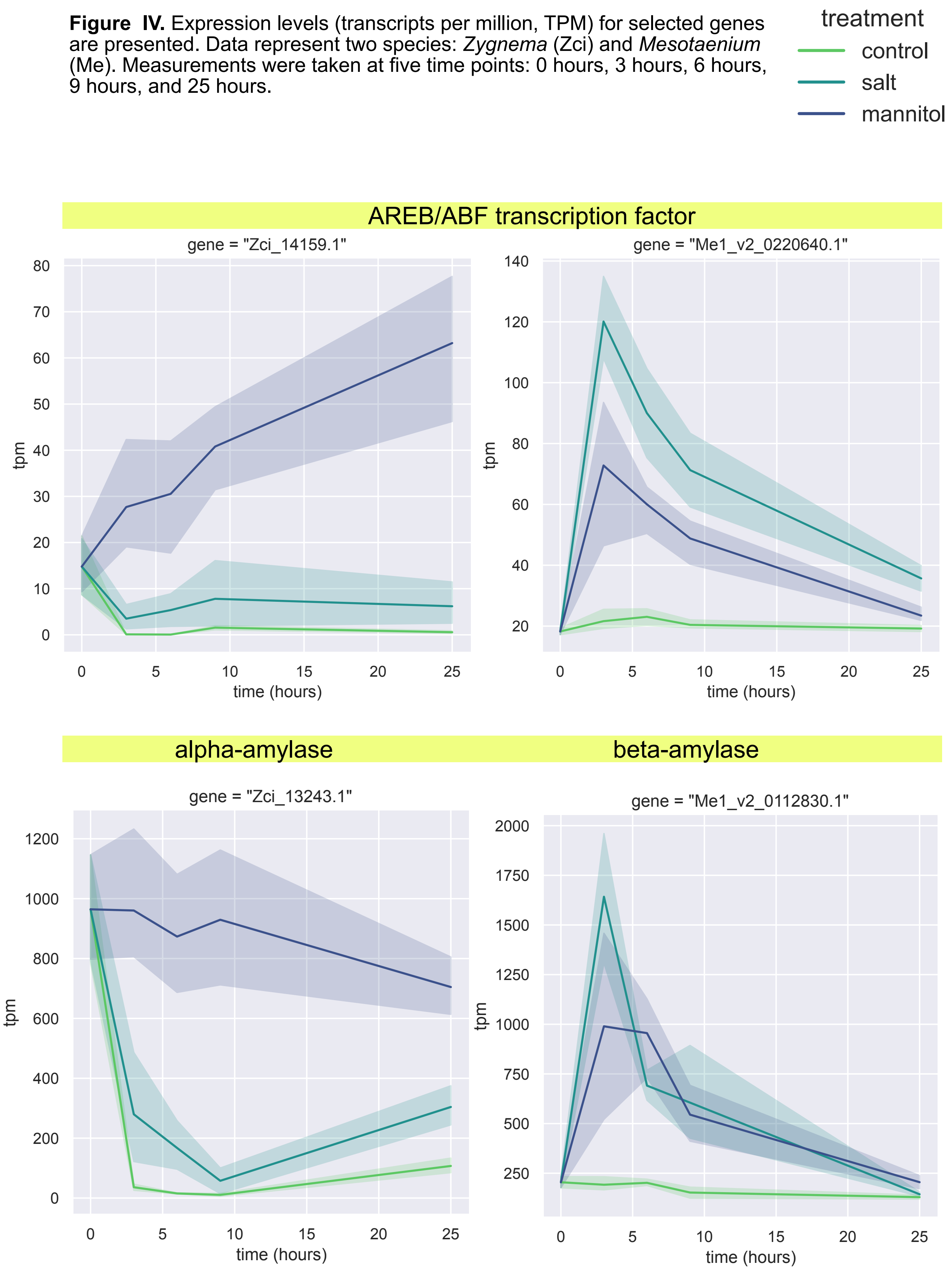

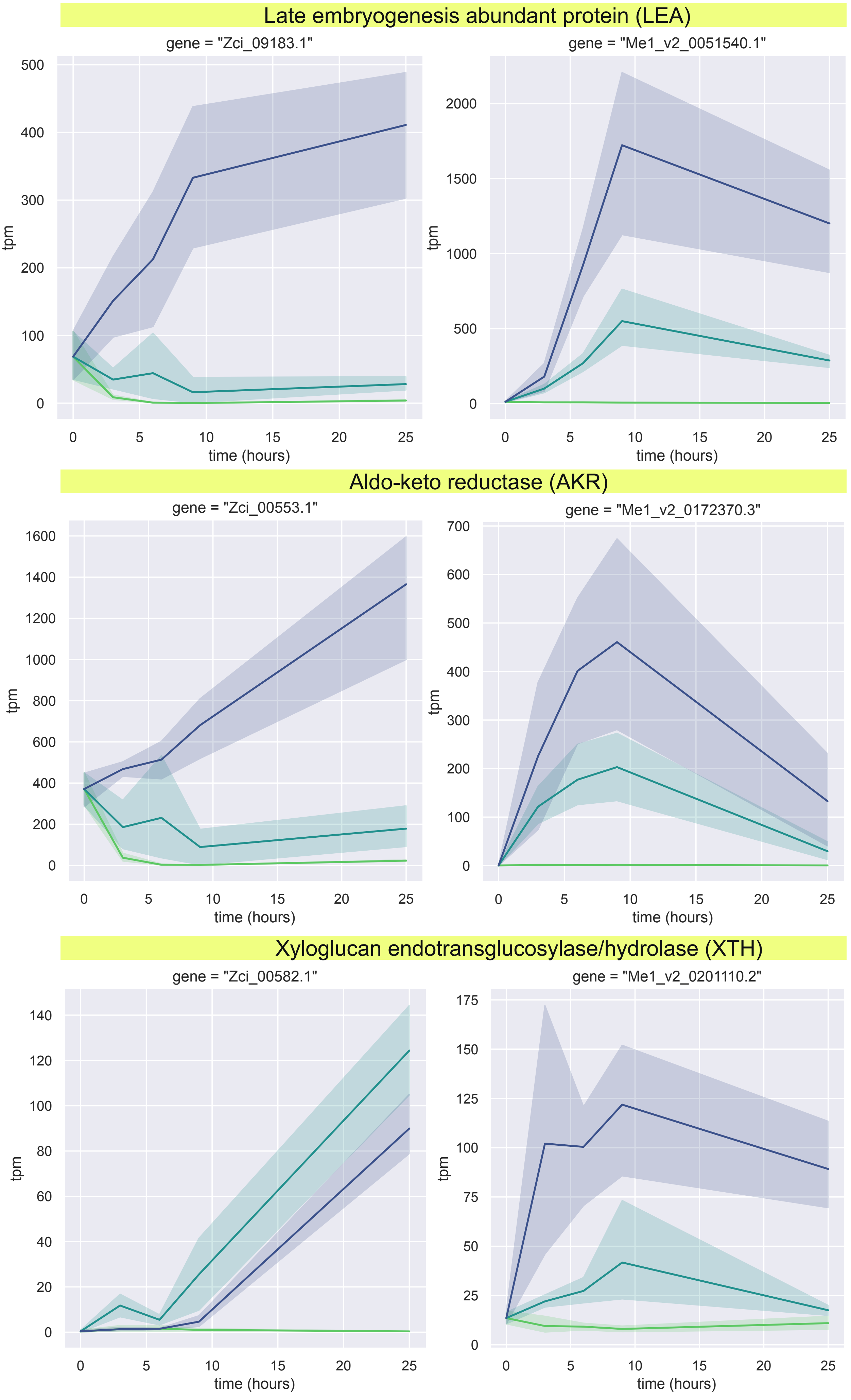

Late embryogenesis abundant protein (LEA)
gene = "Zci_09183.1"
gene = "Me1_v2_0051540.1"
Aldo-keto reductase (AKR)
gene = "Zci_00553.1"
gene = "Me1_v2_0172370.3"
Xyloglucan endotransglucosylase/hydrolase (XTH)
gene = "Zci_00582.1"
gene = "Me1_v2_0201110.2"
tpm
time (hours)

### Fasciclin-Like Arabinogalactan protein (FLA)

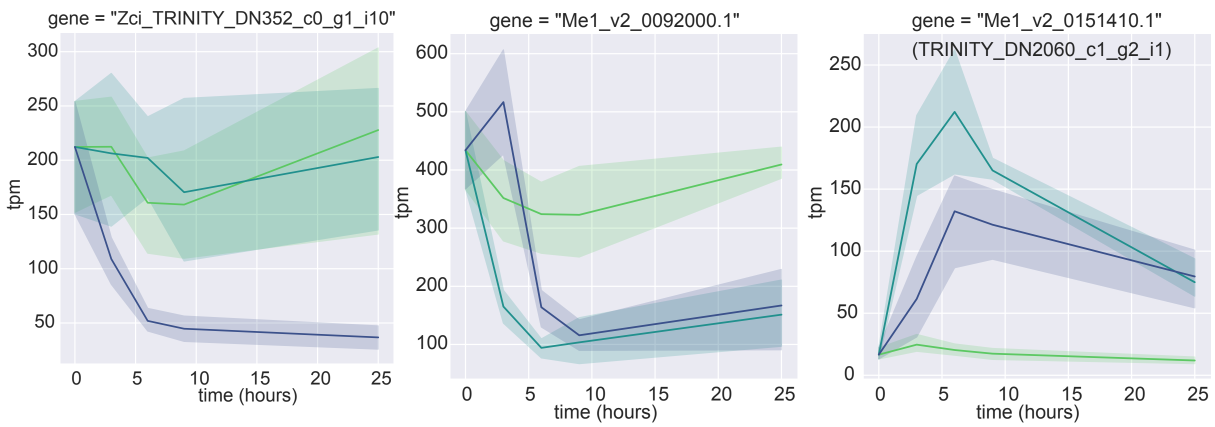


### pectin methylesterase (PME)

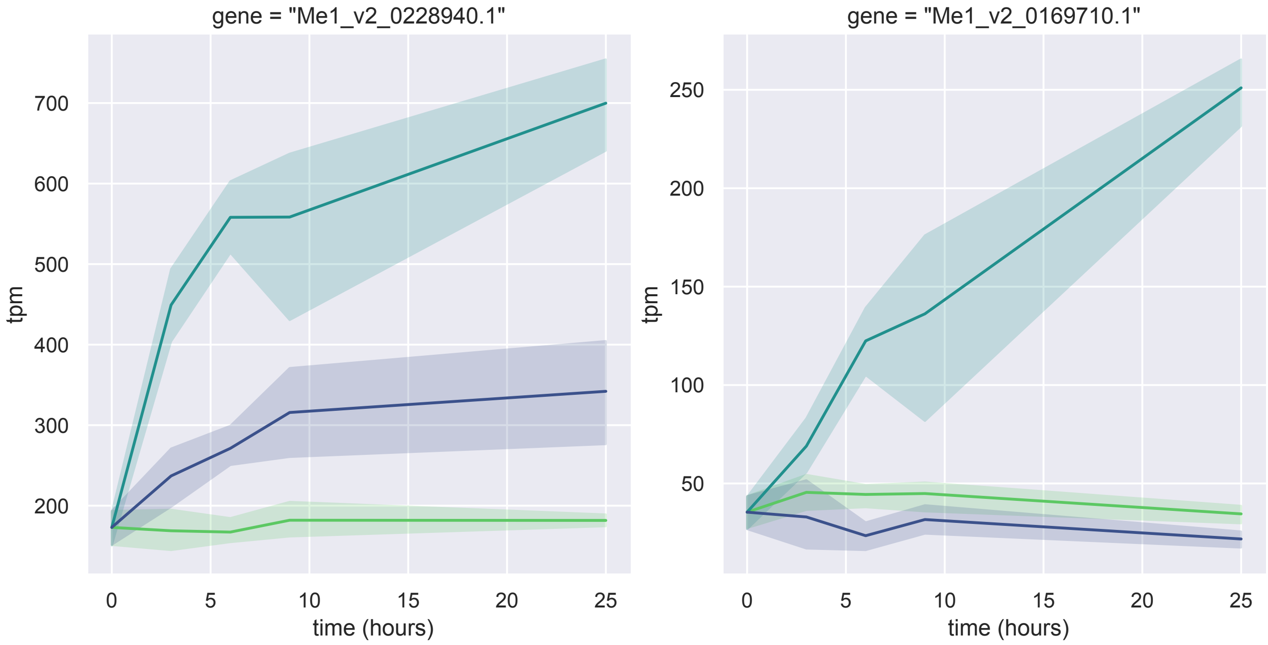
