## Supplementary Figure V for "Systems acclimation to osmotic stress in zygnematophyte cells"

### Supplementary Figure(s) V. Metabolomics

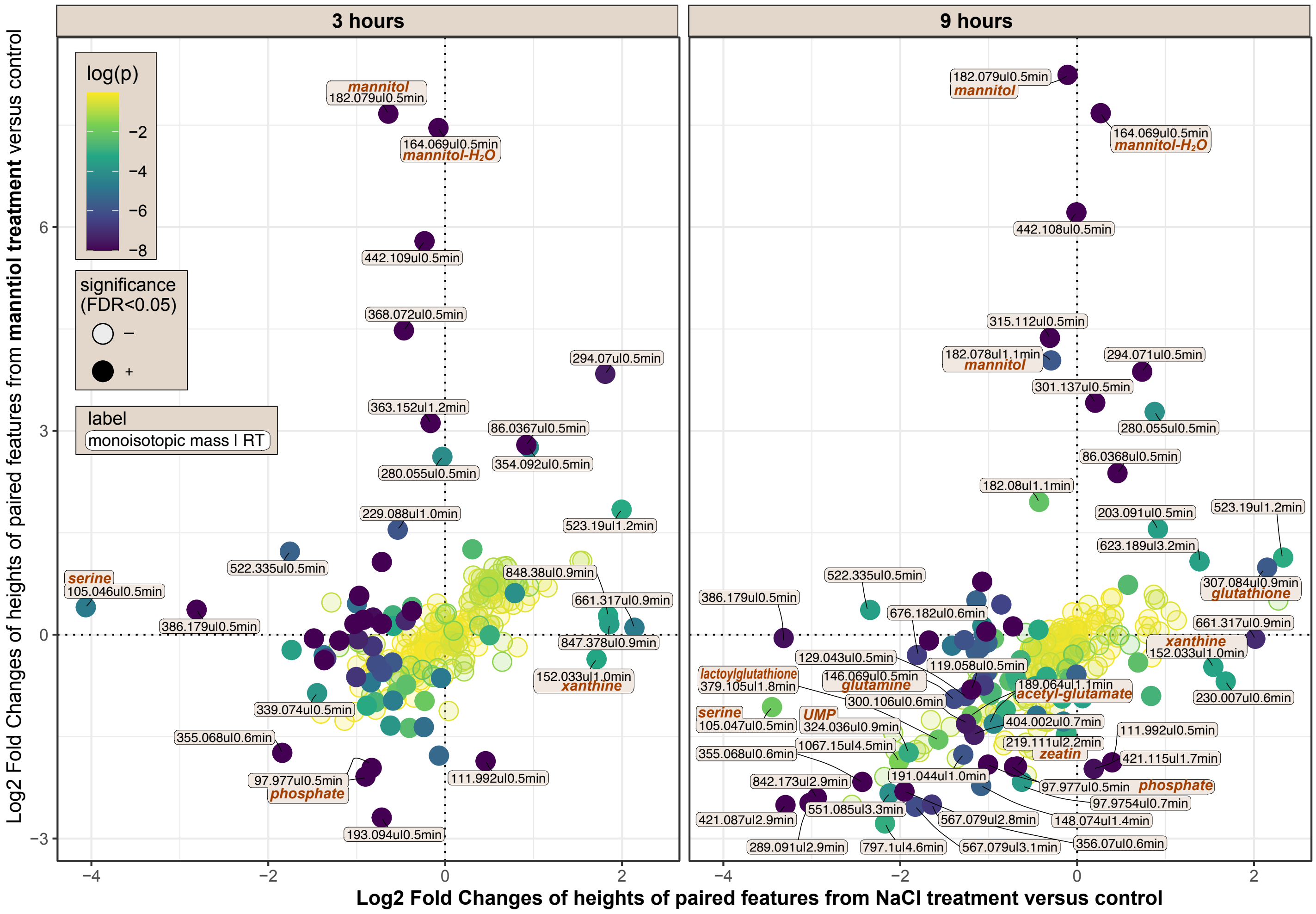


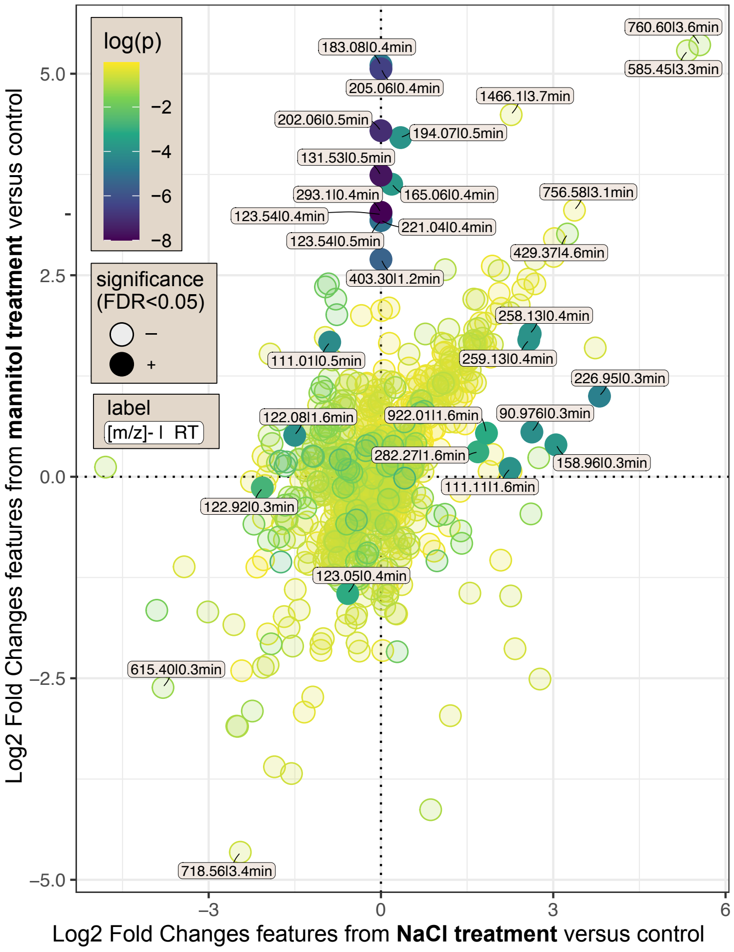


**Figure V.A** Comparison of the Log2 fold changes of features from LC-MS data of metabolite extracts of *Mesotaenium* subjected to osmotic treatments versus control conditions. The osmotic treatments include 0.8 M mannitol and 0.15 M NaCl. The Log2 fold changes for the 0.8 M mannitol treatment are plotted against those for the 0.15 M NaCl treatment.

Top panel) The features shown are detected in both positive and negative ion modes of methanol-based metabolite extracts. Data from the two ion modes were combined for pairs of features where the mass/charge (m/z) difference corresponds to the mass of two hydrogen atoms (2.01568 u), with a maximum allowed deviation of 0.002 u. Additionally, retention times were matched within a maximum deviation of 0.04 minutes. Unpaired features not shown. For paired features with combined absolute Log2 fold changes greater than 2 and a false discovery rate (FDR) less than 0.05, labels indicate the inferred monoisotopic weight (calculated as the average of m/z values in both ion modes) and the retention time. Where applicable, the names of putative compounds are provided based on monoisotopic weights that match entries in the BioCyc databases for *Arabidopsis thaliana*, *Klebsormidium flaccidum*, and *Chlamydomonas reinhardtii*.

Bottom) This panel presents features detected in the positive ion mode of lipid extracts from *Mesotaenium* after 9 hours of treatment. For features with an FDR less than 0.05 and/or a summed Log2 fold change exceeding |6|, labels denote the corresponding m/z values and retention times.

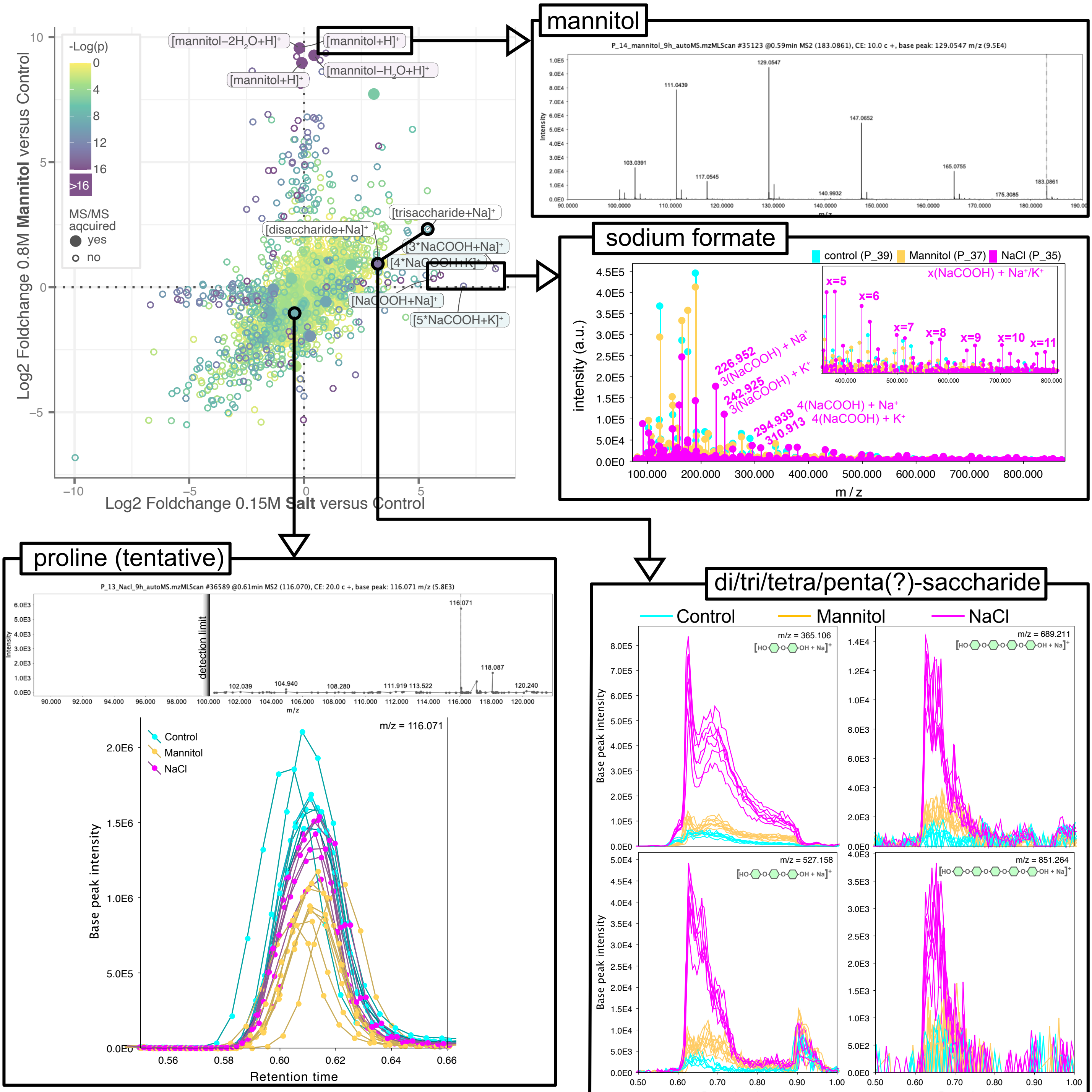


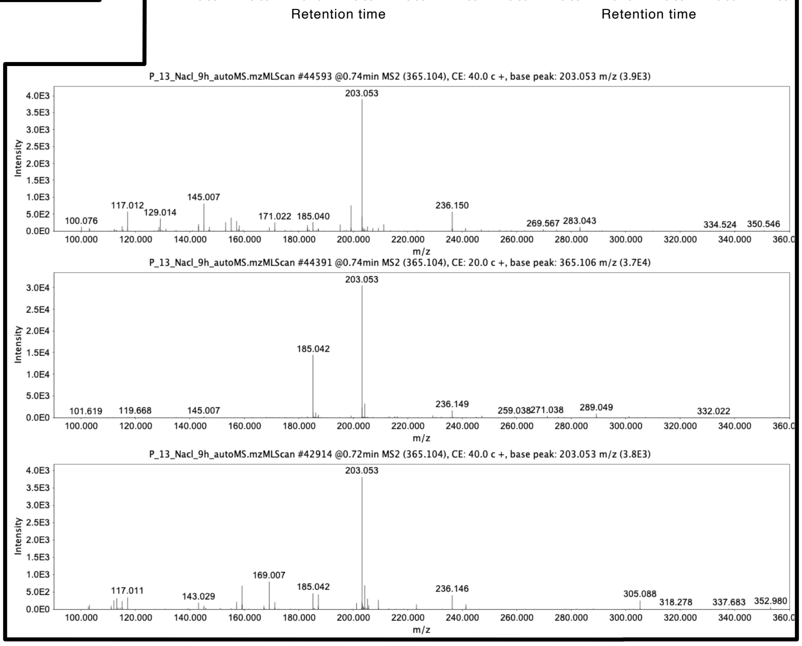

**Figure V.B** Chromatograms and Mass Spectra Supporting Features Visualized in Figure 6a

Mannitol: One of the 33 acquired fragmentation mass spectra tentatively identified as mannitol.

Sodium Formate: Mass spectra from a control sample (scan #28664), an NaCl-treated sample (scan #28636), and a mannitol-treated sample (scan #28638) at a retention time (RT) of 0.49 minutes.

Saccharides: Total ion chromatograms of all samples for features with RTs around 0.64 minutes and m/z values of 365.106, 527.158, 689.211, and 851.264. For the feature at m/z = 365.106, three of the nine acquired fragmentation spectra are shown.

Proline: The feature most likely corresponding to proline (based on m/z) is displayed. The top panel shows the fragmentation spectrum, and the bottom panel shows the total ion chromatograms of all samples.

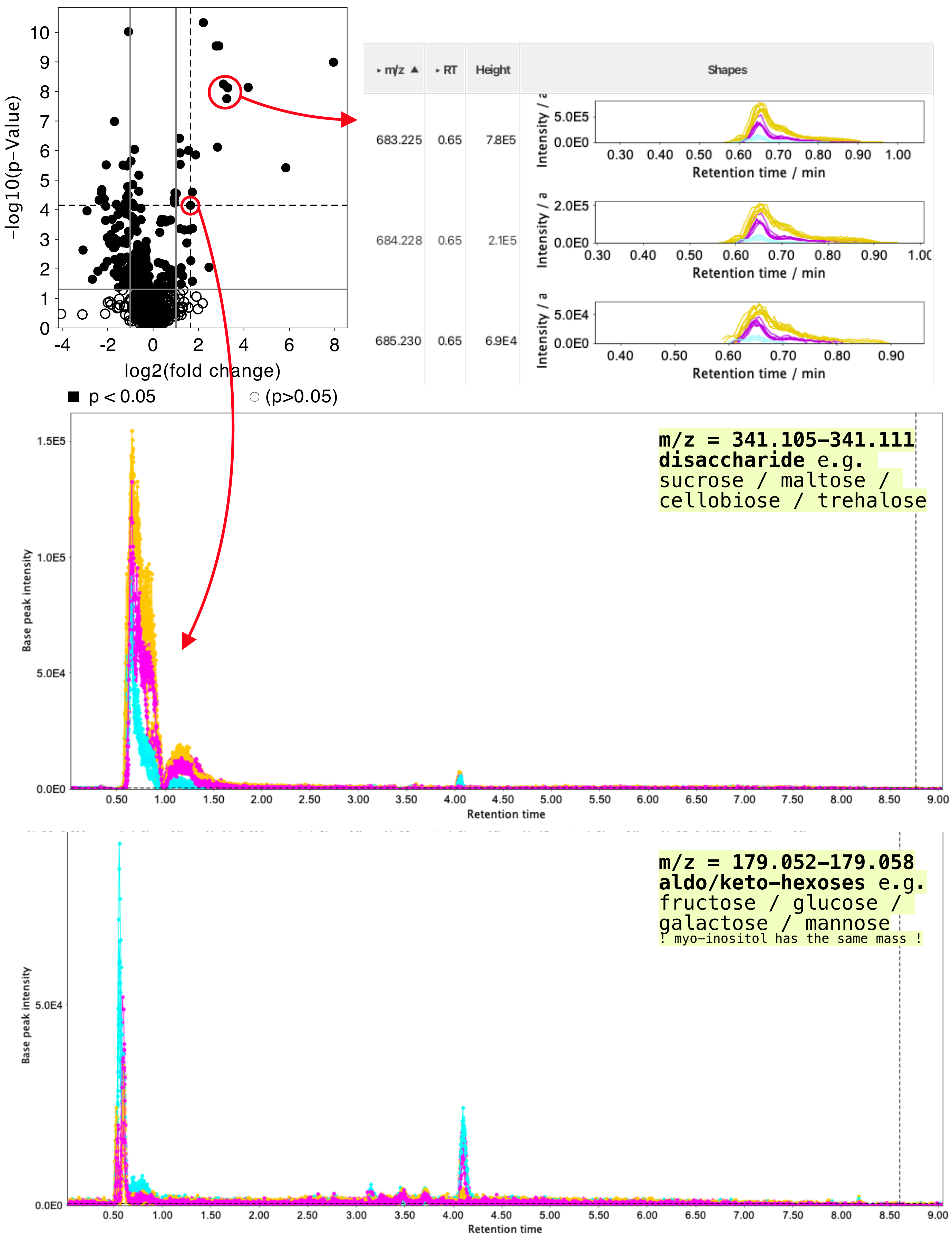


**Figure V.C** Top left panel: Volcano plot illustrating non-gap-filled features in negative ion mode, comparing 9-hour mannitol treatment to 9-hour control treatment of methanol-based metabolite extracts of *Mesotaenium*.
Top right panel: A feature group representing an unidentified compound that exhibits a strong increase in intensity in response to osmotic stress in Mesotaenium.
Bottom panels: Total ion chromatograms of methanol-based extracts from Mesotaenium collected after 9 hours of treatment, analyzed in negative ion mode. The chromatograms highlight two specific m/z ranges: 341.105–341.111, corresponding to the mass of various disaccharides ($[M-H]^-$), and 179.052–179.058, corresponding to the mass of aldo- or keto-hexoses ($[M-H]^-$, e.g., glucose and fructose).
Color coding: NaCl = magenta, Control = cyan, Mannitol = orange.

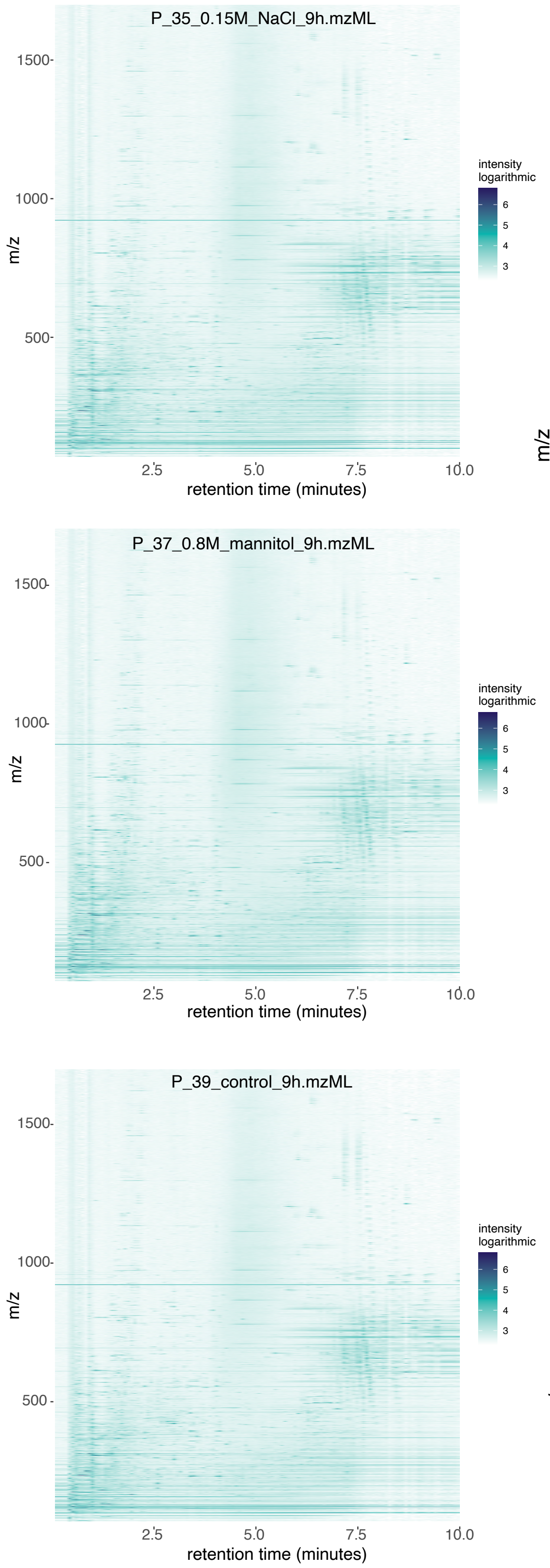


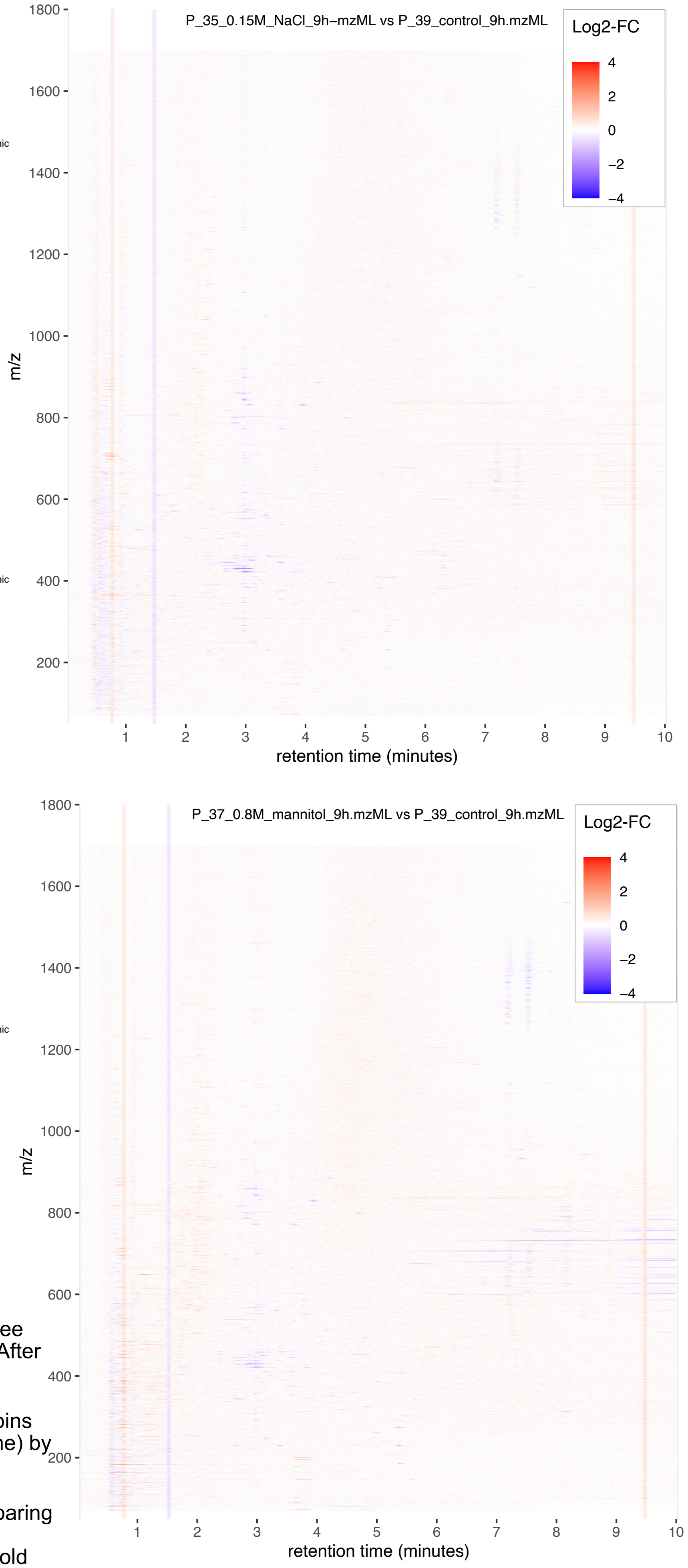


**Figure V.D** 2D LC-MS Chromatograms of three Methanol-Based Extracts from *Mesotaenium* After 9 Hours of Treatment (Positive Ion Mode).

Left Panel: Average signal intensities across bins with dimensions of 0.05 minutes (retention time) by 0.5 (mass-to-charge ratio).

Right Panel: Fold changes of these bins comparing samples treated with 0.15 M NaCl or 0.8 M mannitol to the control sample. For bins with fold changes exceeding |4| (only three bins in total), bins are shown in green.
