## Supplementary Figure VI for "Systems acclimation to osmotic stress in zygnematophyte cells"

### Supplementary Figure VI. Sugar analysis

**A.** Total sugar content

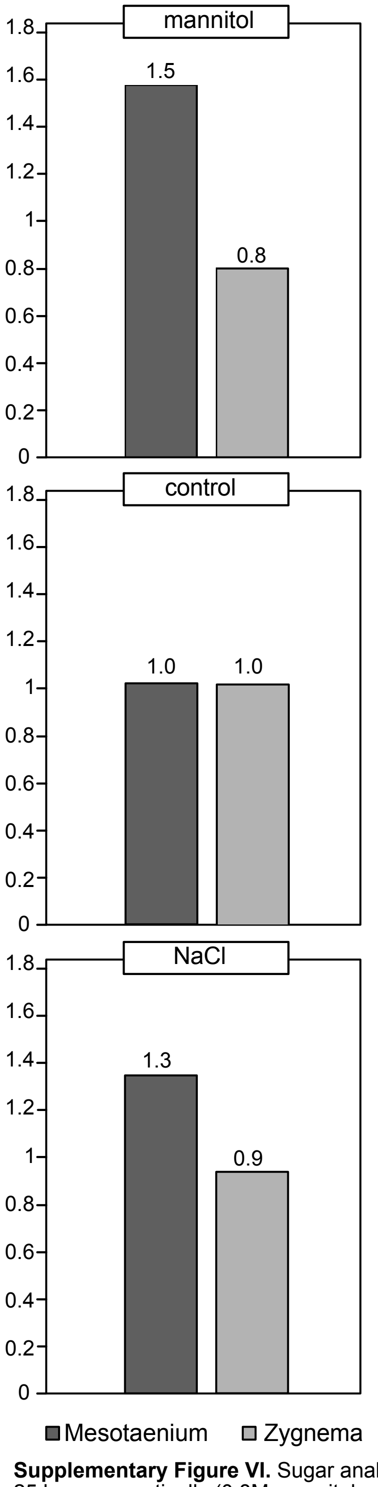


**B.** Uronic acid content

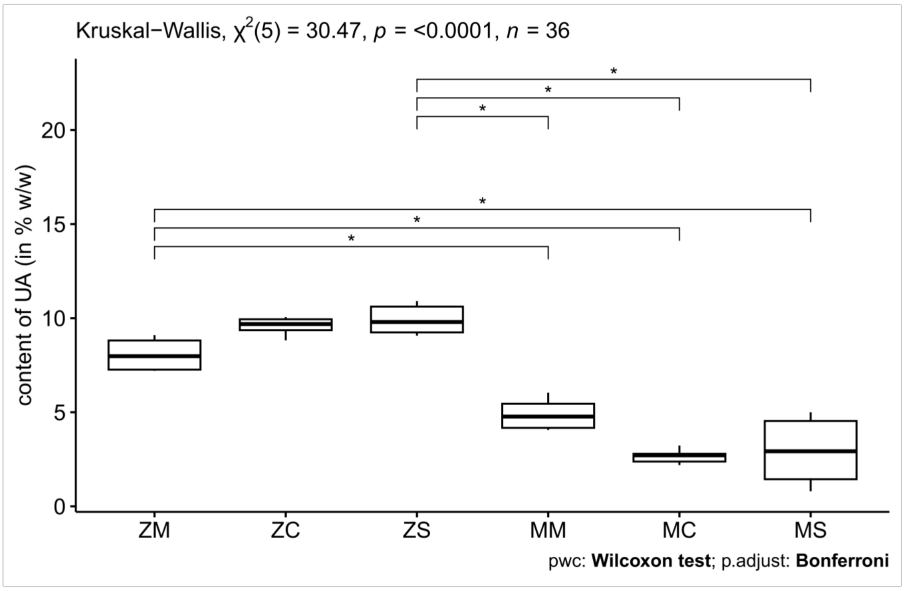


**C.** Hydroxyproline quantification

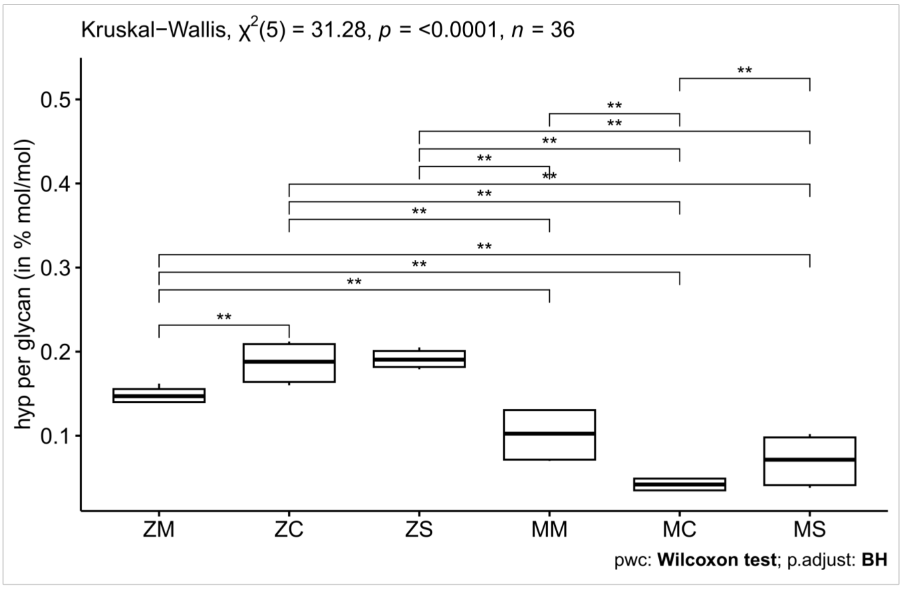


**D.** Neutral sugar composition

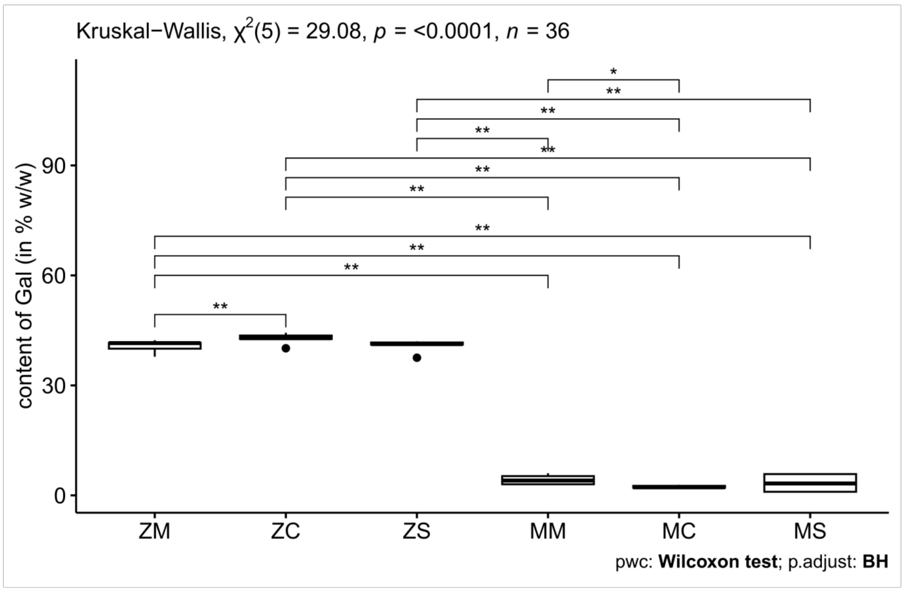


**Supplementary Figure VI.** Sugar analysis of 25 hours osmotically (0.8M mannitol or 0.15M NaCl) stressed algae. Averages shown from 3x technical replecate, 2x biological replicate. **A)** Total sugar content of all neutral and acidic sugars determined by colorimetric assay according to Dubios et al. (1956) **B)** Uronic acid content determined by colorimetric assay according to Blumenkrantz and Asboe-Hansen (1973). **C)** Hydroxyproline colorimentric qauntification according to Stegemann and Stalder (1967). **D)** Neutral sugar composition determined of the molar ratio of the monosaccharides.
ZM: Zygnema - mannitol
ZC: Zygnema - control
ZS: Zygnema - salt
MM: Mesotaenium - mannitol
MC: Mesotaenium - control
MS: Mesotaenium - salt
