## Supplementary Figure VII for "Systems acclimation to osmotic stress in zygnematophyte cells"

### Supplementary Figure(s) VII. β-Yariv reagent staining

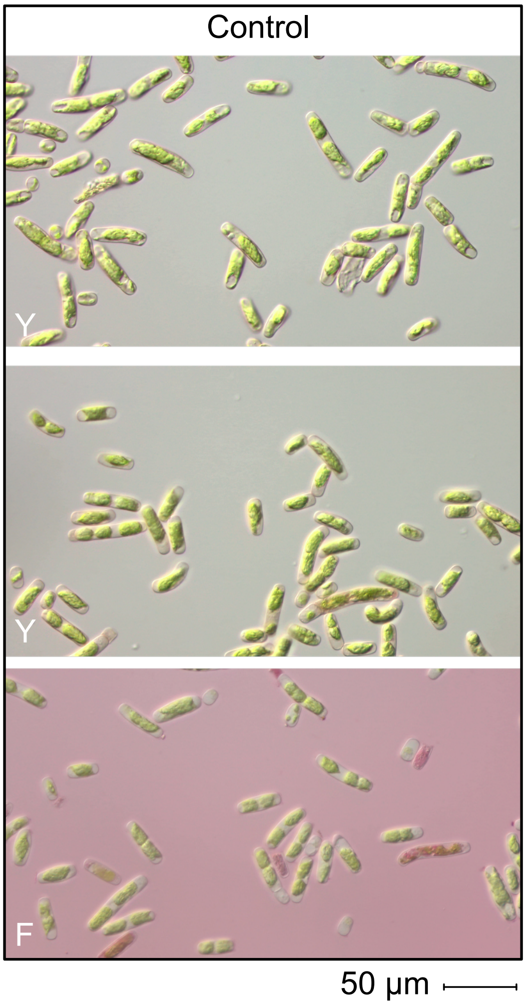


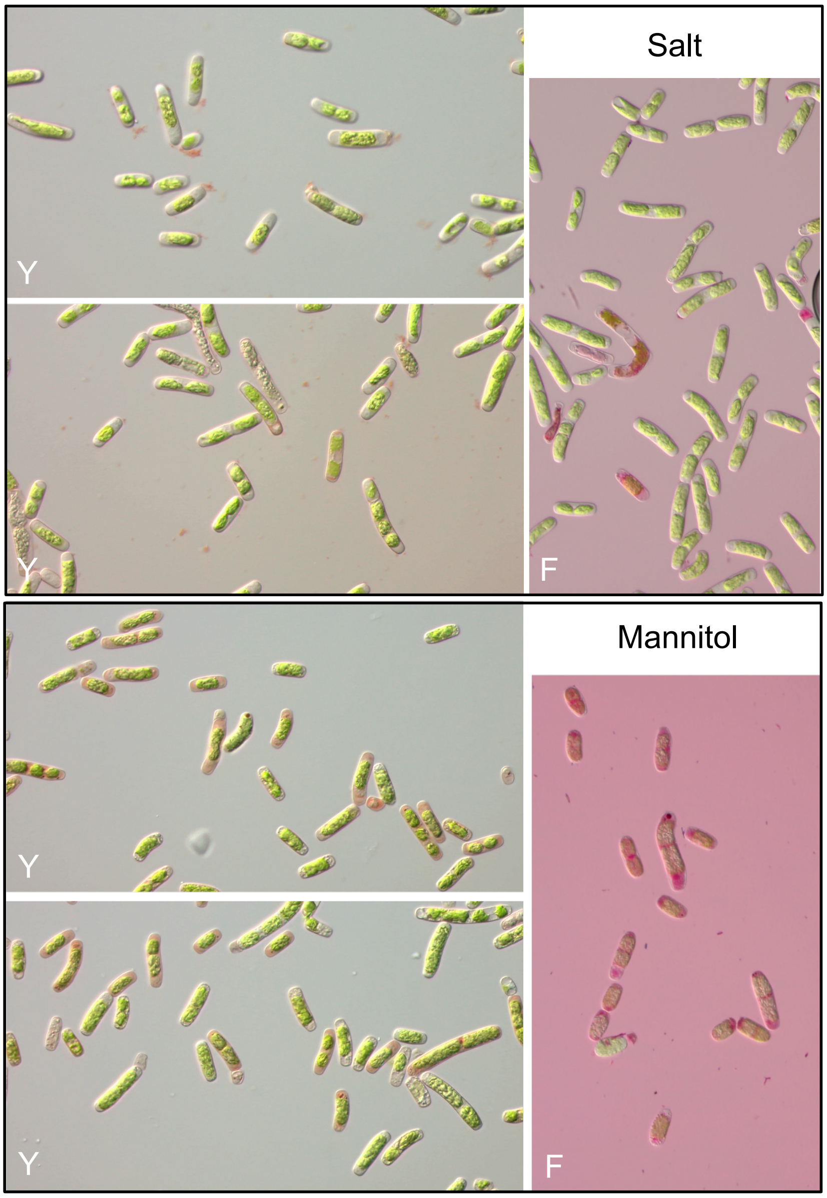


**Figure VII-A**. 200 µM β-yariv (Y) stain or 1% fuchsin (F) with 10 min incubation. Extracellular staining in salt treatments was commonly observed as depicted, ranging from sparse blobs of 1 nm in diameter (bottom) to larger aggregates (top). Such staining was virtually never observed in mannitol treatments, and rarely in control. Staining with 1% fuchsin did not reproduce similar patterns.

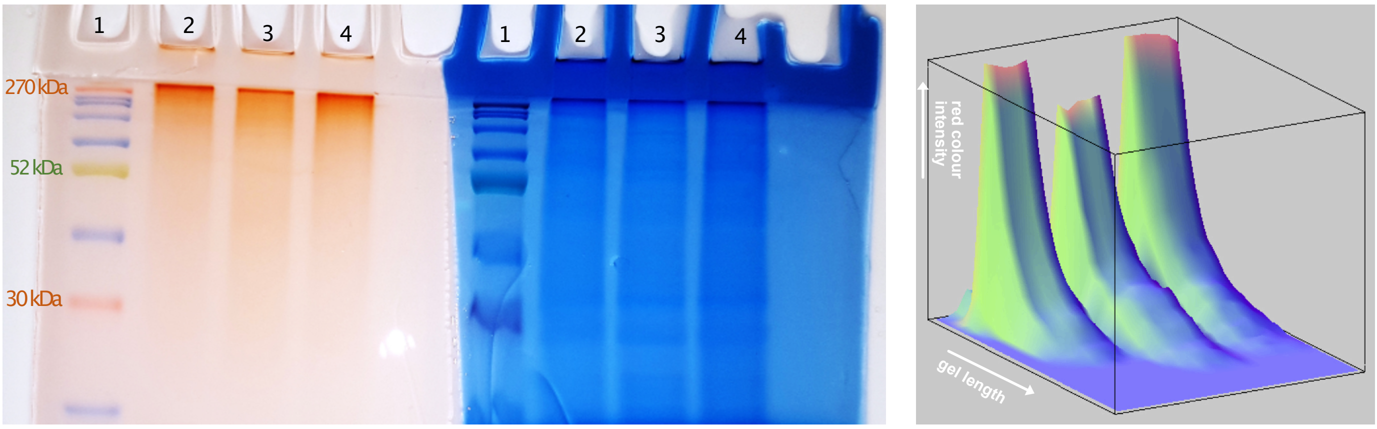


**Figure VII-B.** 12% acrylamide SDS-PAGE gel stained with 0.2% (w/v) ß-yariv reagent on the left side and with Coomassie blue on the right side. Lane 1: abcam prestained protein ladder - extra broad. Lane 2-4: *Mesotaenium* total protein extract, taken from sample after 25 hours of control (2), mannitol (3), or salt (4) treatment. 3D graph on the right show Red channel colour intensities of lanes 2-4 from left to right.
