## Supplementary Figure VIII for "Systems acclimation to osmotic stress in zygnematophyte cells"

### Supplementary Figure(s) VIII. Long term NaCl treatment

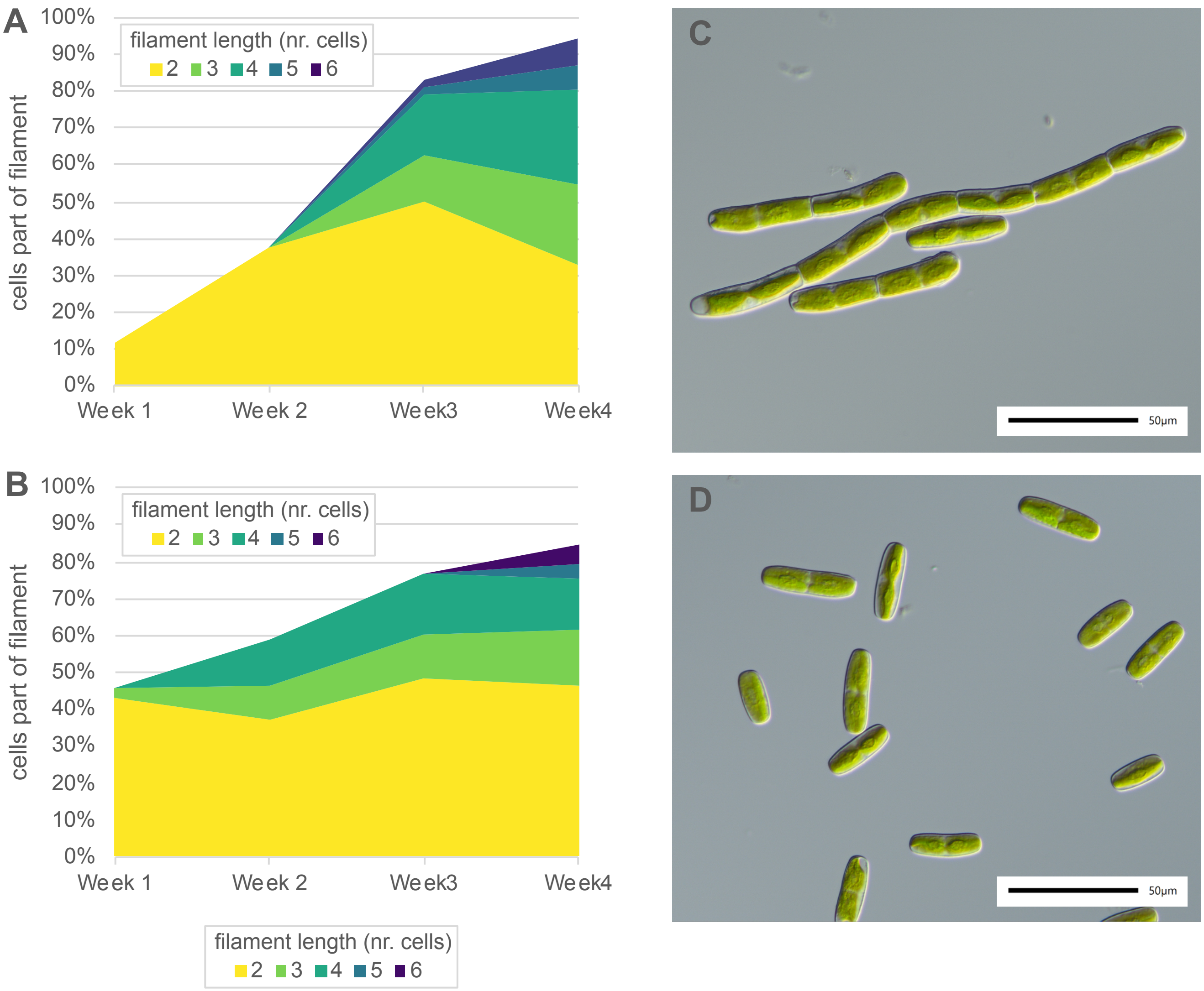


**Figure VIII-A.** NaCl-induced short-filamentous growth of *Mesotaenium endlicherianum* SAG12.97. A) Percentage of cells forming filaments after 1-4 weeks of growth on WHM-agar medium supplemented with 0.15 mM NaCl. No filaments (n≥2) were observed in the control. B) Same as panel A, but cells were grown on top of cellophane. C) Light microscopic image of *M. endlicherianum* after 3 weeks of growth in the presence of 0.15 mM NaCl. D) Light microscopic image of the corresponding NaCl-free control from panel C.

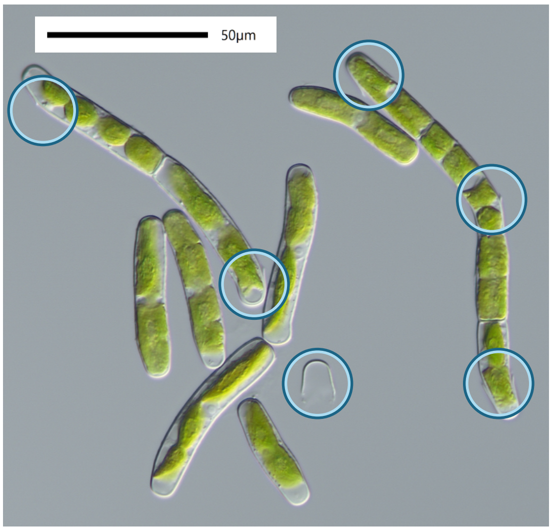


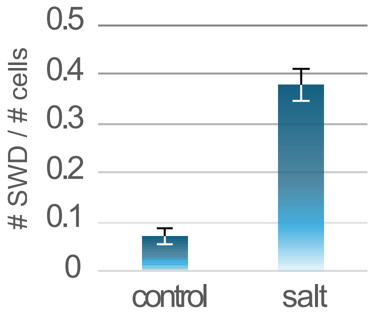


**Figure VIII-B.** Secondary wall deposits/features (SWDs) in *M. endlicherianum* after 2 weeks of growth on cellophane foil over agar-solidified WHM medium supplemented with 0.15 M NaCl. Left: Example of SWDs. Top: Ratio of SWDs to cell count for the treatment with NaCl and the control without NaCl.

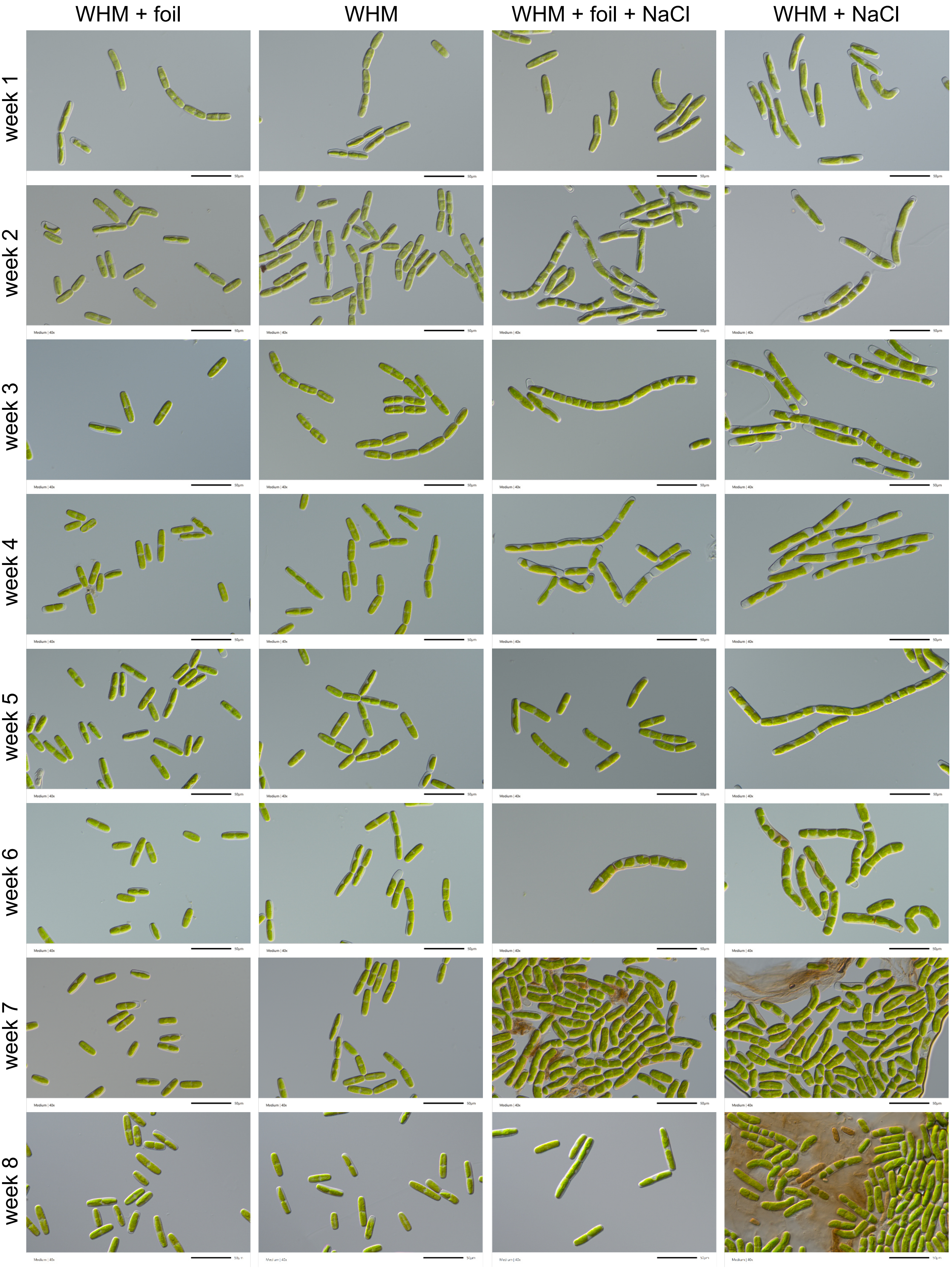


**Figure VIII-C.** Microscopic images showing the growth of *M. endlicherianum* over 8 weeks after inoculation onto agar-solidified WHM medium. The images compare growth conditions with and without a cellophane foil overlay, and with and without the addition of 0.15 M NaCl.
