## Supplementary Table I for "Systems acclimation to osmotic stress in zygnematophyte cells"

### Supplementary Table(s) I. Microscopy

Table I-A. Number of Mesotaenium cells and Zygnema filaments that were analysed per biological replicate (BR) as well as the morphologies they were scanned for.

| | Mesotaenium | | | | | | |
|---|---|---|---|---|---|---|---|
| | BR1 | BR3 | BR4 | BR5 | BR6 | BR7 | Total |
| control | 68 | 208 | 128 | 111 | 291 | 152 | 958 |
| mannitol | 77 | 208 | 292 | 140 | 134 | 194 | 1045 |
| salt | 102 | 215 | 179 | 130 | 177 | 142 | 945 |

| Morphology | Description |
|---|---|
| **Death cell** | Absence of green colour throughout cell. |
| **Broken cells** | Disruption of the cell wall that leads to visible release of cytosol/organelles outwards of the cell. |
| **Cupped cells** | The presence of a small cell-wall enclosed cup on tip of cell that does not contain a visible nucleus and chloroplast. Must still be attached. |
| **Coloured cells** | Presence of orange/brown/red/pink colouration of one or more vescicle within the cell. ("cups" excluded) |
| **Visible tonoplast** | Visible membrane around completely transparant cell compartment(s) (presumably vacuole), that do not reflect light and lack a nucleoles. |
| **LDL cell** | Presence of at least 5 circular droplets of at least 1.5 µm (0.054 in ImageJ) that reflect the light and have no colour |
| **Bend cells** | Bending angle smaller than 160 degrees, cups not included. |
| **plasmolysed** | The plasmamebrane is seperated from the cell wall, resulting in a gap between cell wall and cell contents. |

*Cells were only analsed when sharp and complete on the image. Exception being broken cells, as long as the breakage was clear and cell contained green content.

| | Zygenma | | | | | |
|---|---|---|---|---|---|---|
| | BR1 | BR2 | BR3 | BR4 | BR5 | Total |
| control | 34 | 29 | 32 | 38 | 29 | 162 |
| mannitol | 29 | 35 | 34 | 40 | 31 | 169 |
| salt | 18 | 26 | 33 | 26 | 33 | 136 |

| Morphology | Description |
|---|---|
| **Star-shaped** | At least 1 cell in filament has star-shaped chloroplasts, where at least 1 arm can be distinguished from lobe. |
| **Plasmolysed** | At least 1 cell in filament shows plasmolysis, where there is a visible gap between cell membrane and cell wall. |
| **Broken** | At least 1 cell in filament has a a disrupted cell wall that lead to release of cell content outwards of cell wall. |
| **Bend** | Outer cell wall in filament is bend (curvature < 165 degrees) |

*Filaments were analysed when at least 1 cell was sharp and complete in the image.

Table I-B. Presence or absence of extracellular staining in stress stressments of live SAG 12.97 cells. "+" , Nr. of photos with observed extracellular beta-yariv staining; "-", Nr. of photos with no staining.Treatments denoted as c (control), m (0.8 M mannitol), or s (0.15 NaCl).

| | 100 mM~25 hours | | | | | | 200 mM~25 hours | | | | | | 100 mM~26 hours | | | | | |
|---|---|---|---|---|---|---|---|---|---|---|---|---|---|---|---|---|---|---|
| | + | + | + | - | - | - | + | + | + | - | - | - | + | + | + | - | - | - |
| | c | m | s | c | m | s | c | m | s | c | m | s | c | m | s | c | m | s |
| **BR1A** | 0 | 0 | 0 | 0 | 0 | 0 | 0 | 0 | 4 | 3 | 6 | 0 | 0 | 0 | 0 | 0 | 0 | 0 |
| **BR2A** | 0 | 0 | 0 | 0 | 0 | 0 | 2 | 0 | 8 | 3 | 8 | 0 | 0 | 0 | 0 | 0 | 0 | 0 |
| **BR1B** | 0 | 0 | 4 | 4 | 5 | 0 | 0 | 0 | 0 | 0 | 0 | 0 | 0 | 0 | 0 | 0 | 0 | 0 |
| **BR2B** | 0 | 0 | 3 | 4 | 3 | 0 | 0 | 0 | 0 | 6 | 7 | 6 | 0 | 0 | 0 | 5 | 6 | 4 |
| **BR3B** | 1 | 0 | 0 | 2 | 5 | 4 | 0 | 0 | 0 | 0 | 0 | 0 | 4 | 0 | 6 | 1 | 9 | 2 |
| **BR4B** | 0 | 0 | 4 | 4 | 7 | 0 | 0 | 0 | 5 | 4 | 0 | 3 | 0 | 0 | 7 | 6 | 11 | 0 |
